# Differential Metabolic and Multi-tissue Transcriptomic Responses to Fructose Consumption among Genetically Diverse Mice

**DOI:** 10.1101/439562

**Authors:** Guanglin Zhang, Hyae Ran Byun, Zhe Ying, Montgomery Blencowe, Yuqi Zhao, Jason Hong, Le Shu, Karthick Chella Krishnan, Fernando Gomez-Pinilla, Xia Yang

**Author notes:** **Correspondence:** Xia Yang, Ph.D., Department of Integrative Biology and Physiology, University of California, Los Angeles, Los Angeles, CA 90095, USA, Fernando Gomez-Pinilla, Ph.D., Department of Neurosurgery and Department of Integrative Biology and Physiology, University of California, Los Angeles, Los Angeles, CA 90095, USA.

## Abstract

High fructose intake is a major risk for metabolic syndrome; however, its effects seem to vary across individuals. To determine main factors involved in the inter-individual responses to fructose, we fed inbred mouse strains C57BL/6J (B6), DBA/2J (DBA) and FVB/NJ (FVB) with fructose. DBA mice showed the highest susceptibility to gain adiposity and glucose intolerance. Elevated insulin was found in DBA and FVB mice, and cholesterol levels were uniquely elevated in B6 mice. The transcriptional profiles of liver, hypothalamus, and adipose tissues showed strain- and tissue-specific pathways altered by fructose, such as fatty acid and cholesterol pathways for B6 and PPAR signaling for DBA in liver, and oxidative phosphorylation for B6 and protein processing for DBA in hypothalamus. Using network modeling, we predicted potential strain-specific key regulators of fructose response such as *Fgf21* (DBA) and *Lss* (B6) in liver, and validated strain-biased responses as well as the regulatory actions of *Fgf21* and *Lss* in primary hepatocytes. Our findings support that fructose perturbs individualized tissue networks and pathways and associates with distinct features of metabolic dysfunctions across genetically diverse mice. Our results elucidate the molecular pathways and gene regulatory mechanisms underlying inter-individual variability in response to high fructose diet.

## Introduction

One of the most fascinating questions in biomedical research is why individuals react differently to the same challenge or treatment. In turn, understanding the mechanisms involved in individual variability is crucial for the development of personalized treatments. Metabolic challenges are among the most powerful driving forces of biological adaptations, with long-lasting consequences on homeostatic control and disease stages. The differential phenotypic response to a metabolic perturbation across individuals can stem from differences in the host genome, epigenome, and their transcriptomic expression, which can shed light on the molecular mechanisms underlying inter-individual variability.

High fructose consumption is increasingly recognized as a risk factor for the escalating prevalence of metabolic syndrome (MetS) worldwide, posing significant risks for type 2 diabetes mellitus (T2D), obesity, cardiovascular diseases, and non-alcoholic fatty liver disease (1–5). Previous evidence suggests that fructose elicits different effects on obesity phenotypes across mouse strains (6), but the inter-individual variability in other cardiometabolic phenotypes such as glucose and lipid homeostasis as well as the molecular mechanisms underlying the variability remain unclear. To this end, we systematically examined the metabolic parameters and tissue-specific gene regulation in response to fructose treatment in multiple inbred mouse strains. Here we choose to study C57BL/6J (B6), DBA/2J (DBA), and FVB/NJ (FVB) due to their divergence in genetic composition and variable metabolic responses to diets (6; 7).

To explore the molecular underpinnings of the variable responses to fructose, we focused on transcriptomic studies of hypothalamus, liver, and adipose tissues based on their critical roles in metabolic regulation. The hypothalamus is a critical master regulator of nutrient sensing, food intake, energy expenditure, body weight, and glucose metabolism (8; 9), and is also sensitive to fructose consumption (10). The liver is a main regulator of glucose, steroid, and lipid metabolism (11). White adipose tissue (WAT) is critical for energy and lipid storage, hormone secretion, and inflammation involved in MetS (12; 13). Here we report that fructose has differential effects on the transcriptome of these tissues between mouse strains, which offers insights into key genes and pathways underlying the individualized metabolic perturbations carried by fructose.

## Materials and methods

### Animals and experimental design

Male DBA, B6 and FVB mice (Jackson Laboratory, Bar Harbor, ME) of 8-week age weighing 20-25g were randomly assigned to 8% w/v fructose in drinking water (n=8-12/strain) and control group (n=8-10/strain, drinking water) for 12 weeks. We chose 8% fructose in drinking water to mimic the intake route and the average fructose consumption found in sugar-sweetened beverages (~10% w/v) consumed in humans. All mice had free access to water and a standard Chow diet (Lab Rodent Diet 5001, LabDiet, St Louis, MO) and were maintained under standard housing condition (22-24°C) with 12h light/dark cycle. Daily food and drink intake were monitored on percage basis. Each mouse was examined for changes in a wide spectrum of metabolic phenotypes including body weight, body fat, lean mass, intraperitoneal glucose tolerance test (IPGTT), and serum levels of insulin, glucose, and lipids. Mice were sacrificed at the end of the fructose treatment (12-week), and hypothalamus, liver, and various WAT depots (mesenteric [mWAT], subcutaneous [scWAT], gonadal [gWAT], and intraperitoneal [iWAT]) were dissected out, weighed, flash frozen and stored at −70°C.

### Body weight and body mass composition

Body weight was measured weekly and body mass composition (lean mass, fat mass) was determined by NMR in a Bruker mimispec series mq10 machine (Bruker BioSpin, Freemont, CA) every two weeks.

### IPGTT

Prior to IPGTT at the 1st, 4th, 9th and 12th week of fructose treatment, the animals were fasted overnight at each time point. Each mouse was injected intraperitoneally with 20% glucose at 2g glucose/kg body weight. Blood glucose levels from tail vein were measured at 0, 15, 30, 90, and 120 min after glucose injection using an AlphaTrak portable blood glucose meter (Abbott Laboratories, North Chicago, IL). Area under the curve (AUC) was calculated as a measure of glucose tolerance.

### Serum lipids and glycemic traits

Mice were fasted overnight before sacrifice, and blood samples were collected through retro-orbital bleeding. Serum total cholesterol (TC), high density lipoprotein cholesterol (HDL), un-esterified cholesterol (UC), free fatty acids (FFA), triglycerides (TG), glucose, and insulin were measured by enzymatic colorimetric assays at UCLA GTM Mouse Transfer Core as previously described (10). Low density lipoprotein cholesterol (LDL) was calculated as LDL = TC - HDL - (TG/5).

### RNA sequencing (RNAseq) and data analysis

Total RNA was extracted from a total of 96 hypothalamus, liver, and mWAT tissues (n=6/tissue/group/strain for liver and mWAT; n=4/group/strain for hypothalamus) using All-Prep DNA/RNA/miRNA Universal Kit (Qiagen, CA. USA). mWAT was chosen due to its stronger implications in MetS than other fat depots (14). Sample size was based on previous RNAseq studies in which findings were validated using qPCR and gene perturbation experiments (10; 15; 16). RNA quality was evaluated, and sequencing libraries were prepared and sequenced in pair-end mode on an Illumina HiSeq 4000 sequencing system as previously described (10; 15; 16). To identify the differentially expressed genes (DEGs), we employed a pipeline containing HISAT, StringTie and Ballgown (17) to align reads to mouse genome mm10 (18), assemble transcripts (19), and identify DEGs using a linear model test after filtering genes with low expression levels (FPKM<1) (20). Multiple testing was corrected using the q value approach and false discovery rate (FDR) < 0.05 was used to determine significant DEGs. DEGs were assessed for enrichment of pathways in Gene Ontology and KEGG using the DAVID tool (21; 22). Pathways at FDR < 0.05 were considered significant. RNAseq data was deposited to Gene Expression Omnibus with accession number GSE123896.

### Identification of network key drivers (KDs) of fructose DEGs

To investigate the gene-gene regulations among the fructose DEGs and to identify potential regulators, Bayesian networks of hypothalamus, liver, and adipose tissues were first constructed using an established method (23; 24) (details in Supplementary Methods). The weighted key driver analysis (wKDA) in Mergeomics (25) was employed to predict key regulatory genes, or KDs, of the fructose DEGs from each tissue and each strain. KDs are defined as the network genes whose network neighboring genes were significantly enriched for fructose DEGs based on a Chi-square like statistics followed by FDR assessment in wKDA (Supplementary Methods). Network genes reaching FDR < 0.05 were reported as potential KDs. The gene subnetworks of KDs were visualized using Cytoscape (26).

### Validation of strain-specificity and regulatory role of predicted liver KDs using primary hepatocyte cell cultures

We tested whether two predicted strain-specific liver network KDs, namely *Fgf21* for DBA and *Lss* for B6 indeed show strain-specificity in fructose response. Primary hepatocytes were isolated from male B6 and DBA mice 12 weeks of age (n=3/strain) and subject to direct fructose treatment at 5mM and 45mM concentrations for 6, 12, 24 and 48 hours, with duplicates at each time point/concentration/mouse combination. We then isolated RNA and analyzed expression levels of *Fgf21* and *Lss* using qPCR (Supplementary Methods; Supplementary Table 1). Three-way ANOVA was used to test for the effects of mouse strain, fructose dosage, and time points on *Fgf21* and *Lss* expression. Tukey post-hoc test was used to determine statistical significance in gene expression changes between strains within each concentration and time point.

To validate the role of these predicted KDs in regulating gene subnetworks, we used siRNAs to knockdown *Fgf21* or *Lss*. Six oligos for *Fgf21* and 3 oligos for *Lss* (Sigma-Aldrich, St. Louis, MO; Supplementary Table 1) were tested, among which one for *Fgf21* and two for *Lss* achieved >60% knockdown efficiency. These were used for knockdown experiments, followed by measuring the expression changes in ten predicted downstream network genes using qPCR (Supplementary Methods; Supplementary Table 1).

### Correlation between fructose DEGs and phenotypic characteristics

To assess whether and which of the fructose DEGs were related to the metabolic phenotypes, we calculated the Pearson correlation between the DEGs and individual metabolic traits. Benjamini-Hochberg was used to control FDR.

### Relevance of the fructose DEGs to human GWAS genes of cardiometabolic diseases

Summary statistics of human GWAS for various metabolic phenotypes related to obesity, T2D, and coronary artery disease (CAD) were retrieved from the summary statistics links in GWAS catalog (27) (https://www.ebi.ac.uk/gwas/downloads/summary-statistics; Supplementary Table 2). The fructose DEGs from our mouse study were assessed for enrichment of human GWAS signals using the marker set enrichment analysis (MSEA) in Mergeomics (25) (Supplementary Methods).

### Statistics

For metabolic phenotypes, two-sided Student’s t test was used to determine statistical differences between fructose-fed and control mice within each mouse strain. For phenotypes measured at multiple time points, two-way ANOVA were used to determine the significance of fructose treatment and time points. The statistics of genomic analyses was described in the corresponding sections above.

### Study approval

This study was performed in accordance with National Institutes of Health Guide for the Care and Use of Laboratory Animals. The experimental protocol was approved by the Chancellor’s Animal Research Committee of the University of California at Los Angeles.

## Results

### Distinct metabolic responses to fructose consumption between mouse strains

In response to 12-week fructose consumption, fructose-fed DBA mice gained significant body weight and fat mass compared to the water group, while B6 and FVB showed no differences (Figure 1A-E). In addition, only fructose-fed DBA mice showed a decrease in lean mass (Figure 1F) and increases in the weights of rWAT, mWAT, and scWAT (Figure 1G) at the end of the 12-week fructose treatment. To determine whether caloric intake accounted for these differences, we measured drink and food intake weekly, and found an increasing trend for fructose intake accompanied by a decrease in food intake across the three strains (Supplementary Figure 1). Total caloric intake was similar between treatment groups (Figure 1H). The body composition differences were also not explained by the amount of fructose intake, as B6 (resistant) and DBA (susceptible) had similar fructose intake whereas FVB (resistant) had higher fructose intake (Supplementary Figure 1).

**Figure 1.**
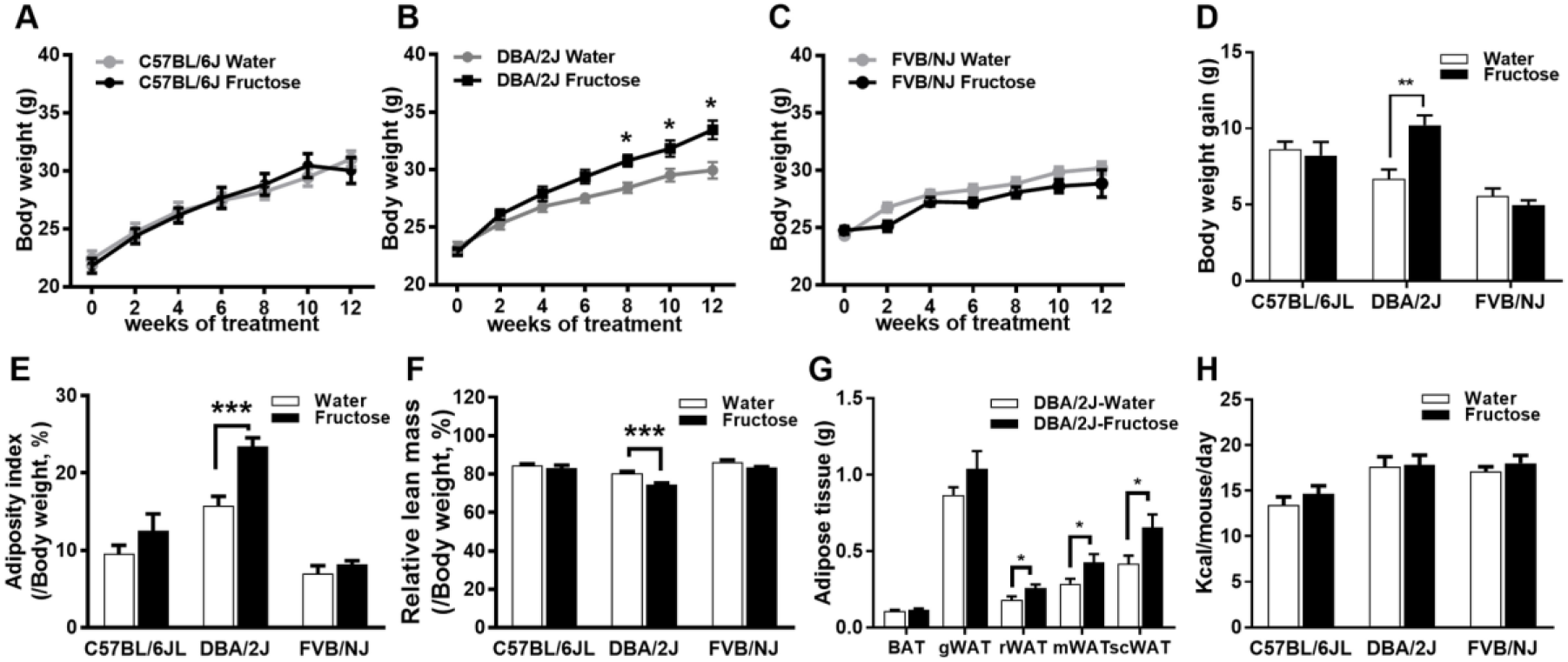
Body weight and body composition changes and caloric intake in three strains of mice in response to fructose consumption. A-C: Cumulative change in body weight in B6 (A), DBA (B) and FVB (C) mice fed normal Chow diet with water or 8% fructose over 12 weeks. Two-way ANNOVA was used to test the differences between treatment groups and time points, followed by Sidak post-hoc analysis to examine treatment effects at individual time points. * denotes *P* < 0.05 from the post-hoc analysis. D: body weight gain at the end of experiment. E-F: Body composition change in three strains of mice, with fat mass (E) and lean mass (F) measured using NMR. G: Individual fat masses of DBA mice. BAT: brown adipose tissue; gWAT: gonadal white adipose tissue; rWAT: retroperitoneal white adipose tissue; mWAT: mesenteric white adipose tissue; scWAT: subcutaneous white adipose tissue. H: Caloric intake of three stains of mice. A-H: Error bars in the graph are standard errors. E-H: * denotes *P* < 0.05 and ** denotes *P* < 0.01 by two-sided Student’s t-test. Sample size n=8-12/group/strain.

As obesity is a risk factor for diabetes and could lead to insulin resistance, we analyzed the glucose tolerance of mice at different time points using IPGTT. Fructose consumption caused increased glucose AUC in DBA mice starting at the 4^th^ week till the 12^th^ week, indicating that fructose impaired glucose homeostasis in DBA. In contrast, B6 and FVB exhibited no difference between treatment groups (Figure 2A-C). No significant difference in serum glucose levels was observed between treatment groups for any of the strains (Fig. 2D), but DBA and FVB showed significantly elevated serum insulin levels in fructose-fed mice (Fig. 2E).

**Figure 2.**
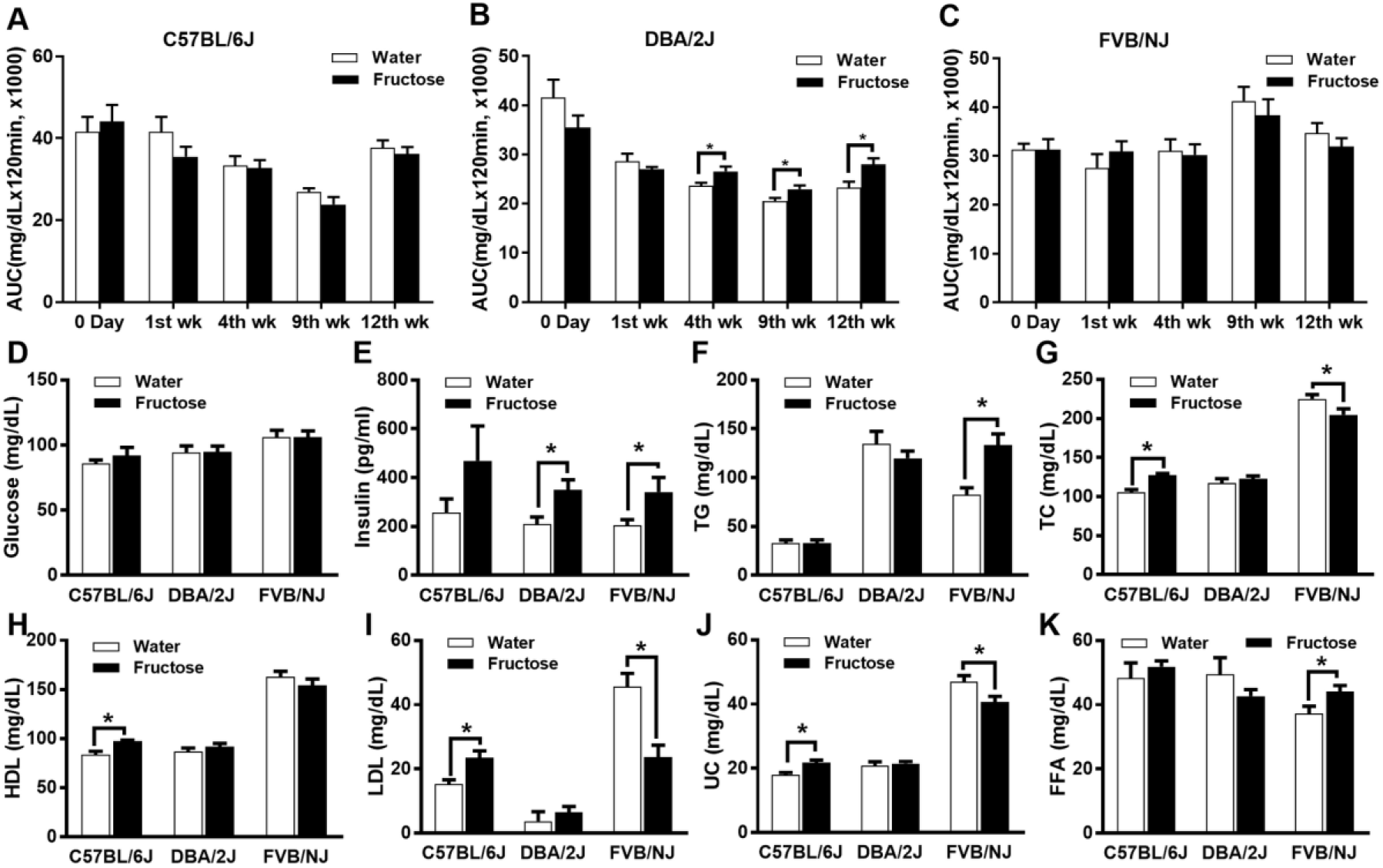
Glucose metabolism and lipid changes in three strains of mice in response to fructose consumption. A-C: Glucose tolerance analysis using IPGTT were conducted at baseline (0 day), 1^st^, 4^th^,9^th^ and 12^th^ week shown as area under the curve (AUC). D: Plasma glucose. E: Plasma insulin. F: Plasma triglyceride. G: Plasma total cholesterol. H: HDL. I: LDL. J: Unesterified cholesterol. K: Free fatty acid levels. A-K: sample size n=8-10 per group. \**P* < 0.05 and \*\**P* < 0.01 by two-sided Student’s t-test.

Lastly, we assessed the influence of fructose consumption on serum lipid profiles. In contrast to the stronger obesity and diabetic phenotypes in fructose-fed DBA mice, lipid traits did not change in DBA. Fructose-fed B6 mice had increased levels of TC, UC, HDL and LDL, while fructose-fed FVB mice displayed elevated TG and FFA but reduced TC, LDL and UC (Figure 2F-K).

Altogether, the above results strongly support that mice with diverse genetic backgrounds exhibit distinct metabolic responses to fructose.

### Distinct transcriptomic changes in response to fructose in metabolic tissues of different mouse strains

To explore the potential mechanisms underlying the differences in metabolic phenotypes among the mouse strains, we examined the transcriptomic alterations in tissues relevant to nutrient sensing, energy homeostasis, and metabolic regulation. RNAseq of tissues from B6, DBA and FVB mice identified 578, 760 and 246 DEGs in liver, and 157, 43 and 9 DEGs in hypothalamus, and 1043, 884, and 427 DEGs in the adipose tissues, respectively, at a threshold of FDR < 0.05 (Figure 3A-C; top DEGs in Table 1; full DEGs in Supplementary Table 3). Based on the DEG numbers, DBA liver and adipose tissues and all three tissues from B6 appear to be more sensitive to fructose compared to FVB.

**Figure 3.**
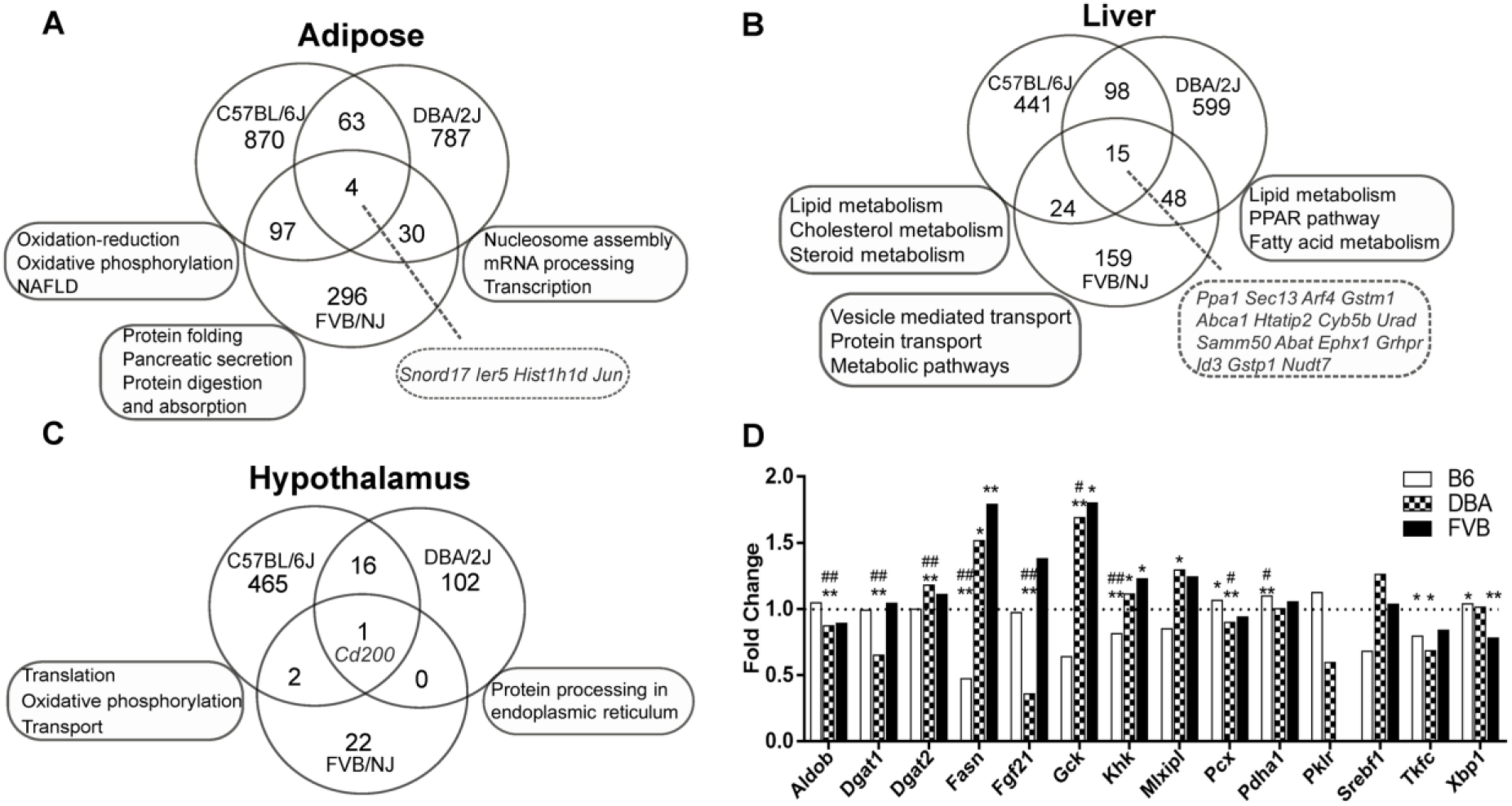
Differentially expressed genes (DEGs) and representative over-represented pathways in adipose, liver and hypothalamus of three mouse strains. A-C: Venn diagrams of adipose (A), liver (B), and hypothalamus (C) DEGs and pathways across three strains. D: Fold change and significance of gene expression differences between fructose and control groups for known target genes of fructose, ChREBP (encoded by *Mlxip1*), SREBP-1c (encoded by *Srebf1*) in the three mouse strains. Fold change for each gene is the normalized average expression level in fructose mice over that of the control mice. * *P* < 0.05, ** *P* < 0.01, # FDR<5%, ## FDR<1% in RNAseq analysis using a linear model. Sample size n=6/group/tissue/strain for liver and adipose tissues; n=4/group/strain for hypothalamus tissue.

**Table 1.**
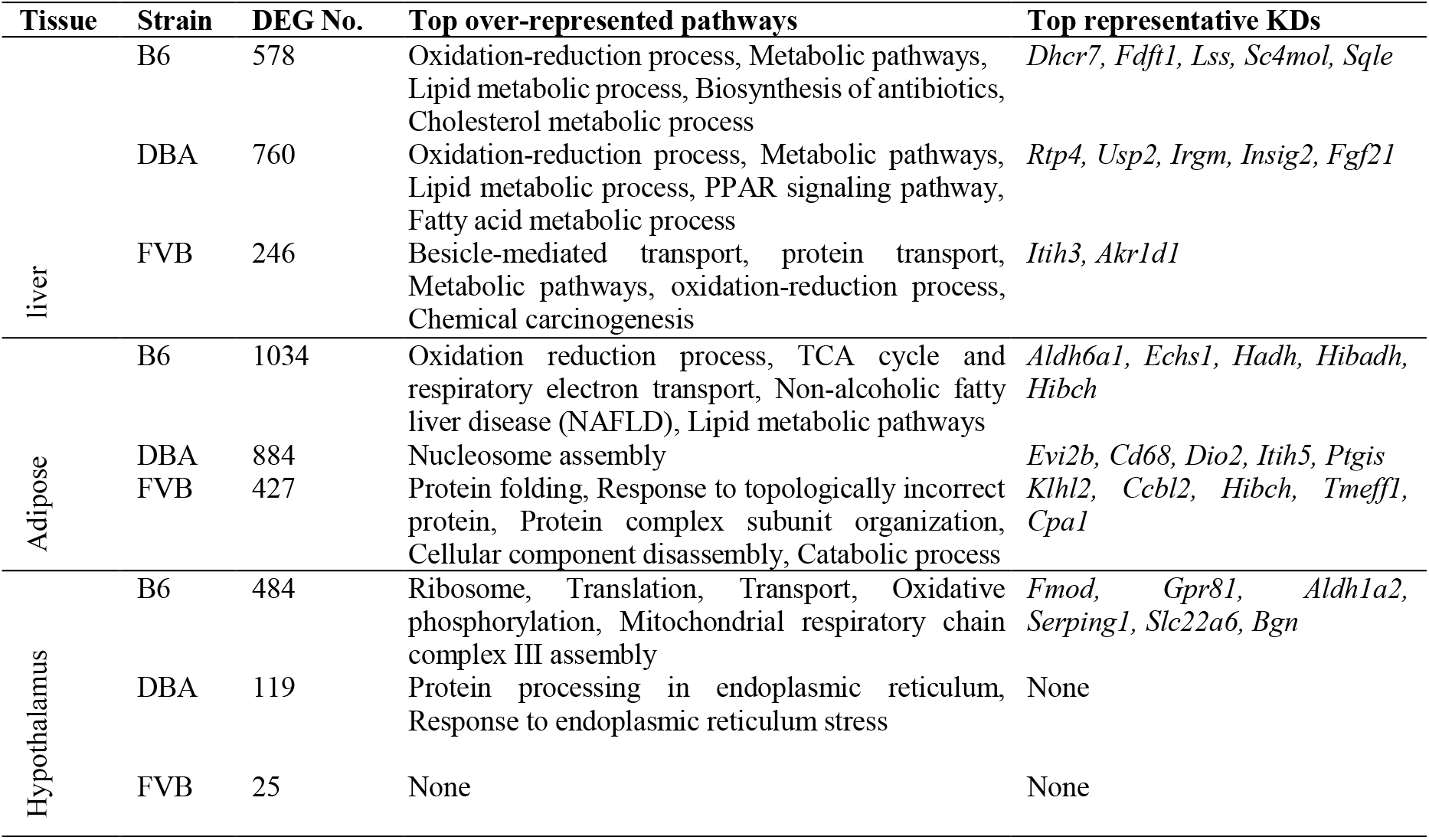
Summary of DEGs in adipose, liver and hypothalamus tissues of three mouse strains at FDR< 0.05 and their top representative key drivers (KDs). Full lists of DEGs, pathways, and KDs are in Supplementary Tables 3-5.

Comparison of the DEGs within each tissue across mouse strains revealed high strain-specificity (Figure 3A-C). Overall, only 4 adipose DEGs (*Snord17, Ier5, Histlhld, Jun*), 15 liver DEGs (*Ppa1, Sec13, Arf4, Gstm1, Abca1, Htatip2, Gstp1, Cyb5b, Abat, Samm50, Urad, Ephx1, Grhpr, Nudt7, Id3*) and one hypothalamus DEG (*Cd200*) overlapped across strains (Figure 3 A-C). Among these, only three genes (*Jun* from adipose tissue, *Id3* from liver, and *Cd200* from hypothalamus) showed consistent direction of changes across strains (Supplementary Table 3).

Previous studies have revealed ChREBP and SREBP-1c as liver transcription factors that regulates fructose activities in liver (28–30). We examined the expression of ChREBP (gene *Mlxip1*), SREBP-1c (gene *Srebf1*), and their known target genes in our liver transcriptome data. *Mlxip1* was significantly altered only in DBA while no changes in *Srebf1* were found in any of the strains. However, many of their target genes such as *Aldob, Dgat1, Dgat2, Fgf21* and *Gck* were significantly altered by fructose in liver, particularly in DBA mice (Figure 3D). Of the genes involved in fructose metabolism (29; 31), *Khk* was downregulated in B6 while upregulated in DBA and FVB, and *Aldob* was downregulated in DBA (Figure 3D).

### Functional categorization of fructose DEGs reveals alterations of strain- and tissue-specific pathways

We annotated the functions of the DEGs and identified tissue- and strain-specific biological pathways affected by fructose (top pathways in Table 1; full results in Supplementary Table 4). Within adipose tissue, B6 DEGs were mainly related to oxidation pathways, DBA DEGs were enriched for nucleosome assembly genes, and protein folding/processing pathways were enriched in FVB DEGs. For liver tissue, B6 DEGs were over-represented with oxidation, lipid and cholesterol metabolic pathways, DBA DEGs were related to oxidation, PPAR signaling, and fatty acid, lipid and cholesterol metabolic pathways, and FVB DEGs were enriched for genes involved in vesicle mediated transport. With respect to the hypothalamus, B6 DEGs were involved in oxidative phosphorylation and protein translation while DBA DEGs were related to protein processing.

### wKDA analysis prioritized strain- and tissue-specific key drivers (KDs) of fructose DEGs

To explore the potential regulatory genes upstream of the fructose-induced DEGs, we used a data-driven network analysis, wKDA, to pinpoint key regulatory genes of fructose activities in individual tissues/strains. This type of analysis previously identified novel regulators of various diseases which were subsequently validated via gene perturbation experiments (10; 32–34). Using wKDA, we predicted KDs of the fructose DEGs from each tissue/strain using tissue-specific gene regulatory networks (top KDs in Table 1; full lists in Supplementary Table 5).

In liver, the top KDs for B6 DEGs were mostly related to sterol biosynthesis such as *Dhcr7, Fdft1*, and *Lss*. The top KDs for DBA included *Fgf21*, a target of ChREBP and a metabolic regulator for insulin sensitivity and obesity (6; 35), *Insig2*, a negative regulator of SREBP targeting lipid metabolism (36), *Irgm*, which induces inflammatory cytokine production (37), and *Arntl*, a key circadian rhythm gene. FVB top liver KDs included *Itih3*, implicated in liver glutathione metabolism (38), and *Akr1d1*, important for bile acid and steroid hormone metabolism (39). For hypothalamus, KDs were identified only for the B6 DEGs, including genes involved in extracellular matrix (e.g., *Fmod, Foxc2, Bgn*). For adipose tissue, DBA-specific KDs are involved in inflammation (*Cd68, Itgis*), obesity (*Dio2*), and cell cycle (*Evi2b*, *Ptpn18*); B6-specific KDs are related to branched chain amino acid (BCAA) metabolism (*Aldh6a1, Echs1, Hadh, Hibadh, Hibch*); FVB KDs are related to kynurenine pathway which is activated in obesity (40) (*Ccbl2*) and BCAA metabolism (*Hibch, Cpa1*). As shown in Figure 4A-C, the top KDs in each tissue orchestrate numerous strain-specific DEGs and there were distinct subnetworks highlighting unique organization of DEGs from each strain.

**Figure 4.**
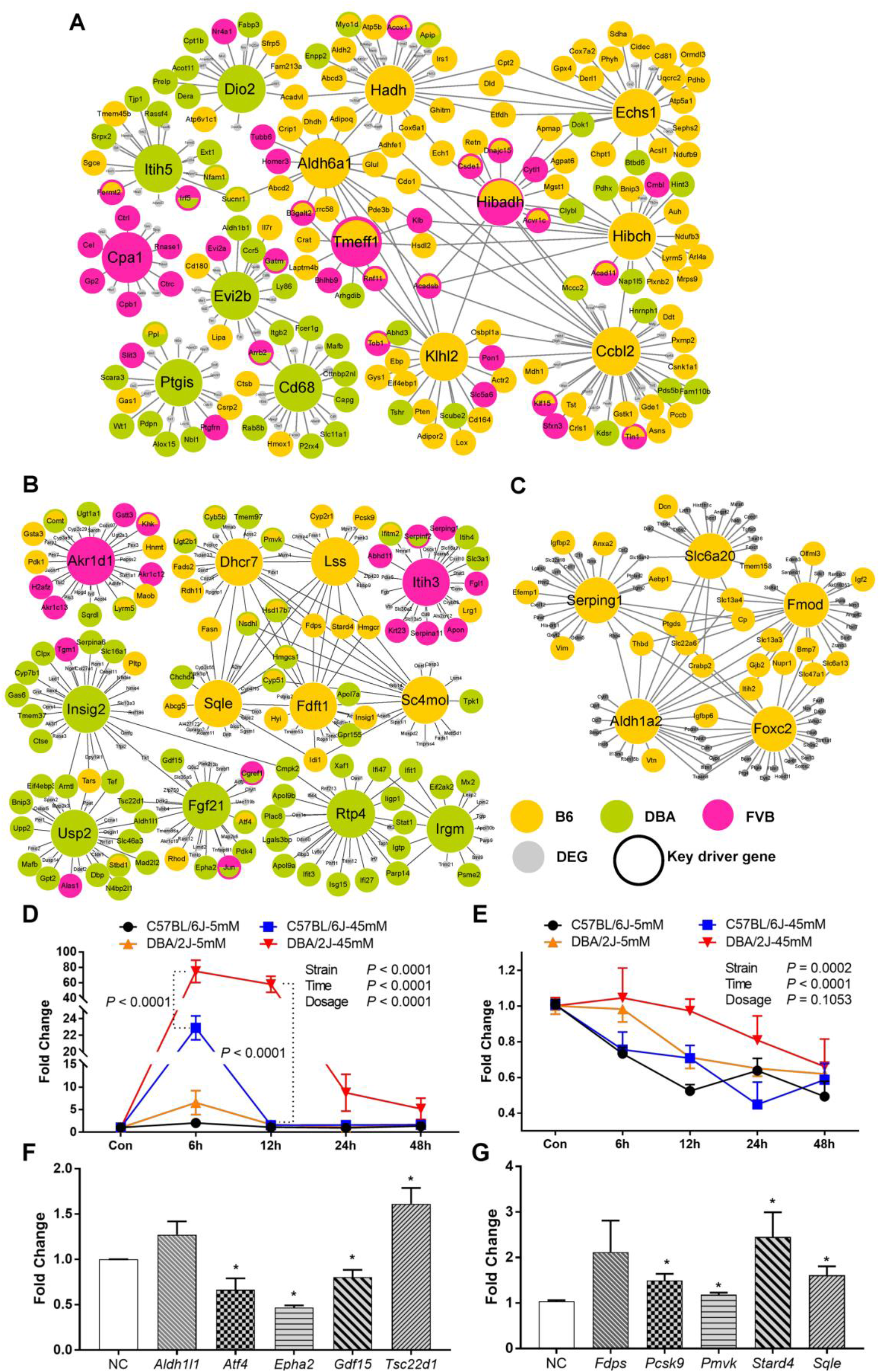
Gene subnetworks and top network key drivers (KDs) of DEGs in three strains of mice and validation of select strain-specific liver KDs in primary hepatocytes. A: Top adipose KDs and subnetworks. B: Top liver KDs and subnetworks. C: Top hypothalamus KDs and subnetworks. D: Direct response to fructose for DBA-specific KD *Fgf21* is stronger in DBA hepatocytes than in B6 hepatocytes. E: Direct response to fructose for B6-specific KD *Lss* is stronger in B6 hepatocytes than in DBA hepatocytes. D-E: P values for strain, fructose concentration, and time points are from 3-way ANOVA. At each concentration and time point, p values between mouse strains are from Tukey post-hoc analysis. Sample size n=3 (2 replicates/mice) per strain per concentration per time point. F: Expression changes of genes in *Fgf21-driven* network after siRNA-mediated knockdown of *Fgf21* in DBA primary hepatocytes. Genes surrounding *Fgf21* were selected from Figure 4B. G: Expression changes of genes in *Lss*-driven network after siRNA-mediated knockdown of *Lss* in B6 primary hepatocytes. Genes surrounding *Lss* were selected from Figure 4B. F-G: NC represents scrambled siRNAs as negative controls. \**P* < 0.05 and \*\**P* < 0.01 by two-sided Student’s t-test. Sample size n=3 (2 repli cates/siRNA).

### Experimental validation of strain- and tissue-specific KDs

*Fmod* and *Bgn* are both B6-specific hypothalamic KDs and were previously found to be hypothalamic KDs in a rat fructose study (10). Ablation of *Fmod* or *Bgn* in mice led to altered metabolic phenotypes and tissue-specific pathways that overlap with those affected by fructose (10; 41). To validate additional strain-specific KDs, we tested the B6-specific liver KD *Lss* and the DBA-specific liver KD *Fgf21* by treating primary hepatocytes isolated from B6 and DBA with fructose. Agreeing with the strain specificity of the KD predictions based on our liver transcriptome data, *Fgf21* was particularly responsive to fructose in DBA hepatocytes (Figure 4D) and *Lss* was more responsive to fructose in B6 hepatocytes (Figure 4E). Perturbing these predicted KDs using siRNAs also affected liver network genes (selected from Figure 4B) surrounding the respective KDs (Figure 4F-G). Altogether, these results support the importance of the network analysis in predicting functional aspects of gene regulation in response to fructose.

### Relationship between DEGs and metabolic phenotypes in mouse and human

To investigate the relationships between fructose DEGs and cardiometabolic phenotypes, we performed a correlation analysis to identify DEGs that show correlation with individual phenotypes in the three mouse strains. At an unadjusted *P* < 0.01, DBA had the largest numbers of adiposity-correlated DEGs from each of the three tissues (Figure 5A; Supplementary Table 6). In contrast, B6 had the largest number of adipose DEGs correlating with TC and FVB liver DEGs showed correlations with LDL, which are consistent with the observed cholesterol phenotypes in these strains. Few of the hypothalamic DEGs correlated with the metabolic phenotypes measured. At the stringent Benjamini-Hochberg adjusted FDR < 0.05, DBA liver DEGs involved in lipid metabolism and adipogenesis (*St6gal1, Scd1*, and *Pla2g6*) were correlated with adiposity (Fig. 5B-D) and FVB liver DEGs (*Aldh2, Enho, Svip*) were correlated with LDL (Fig. 5E-G).

**Figure 5.**
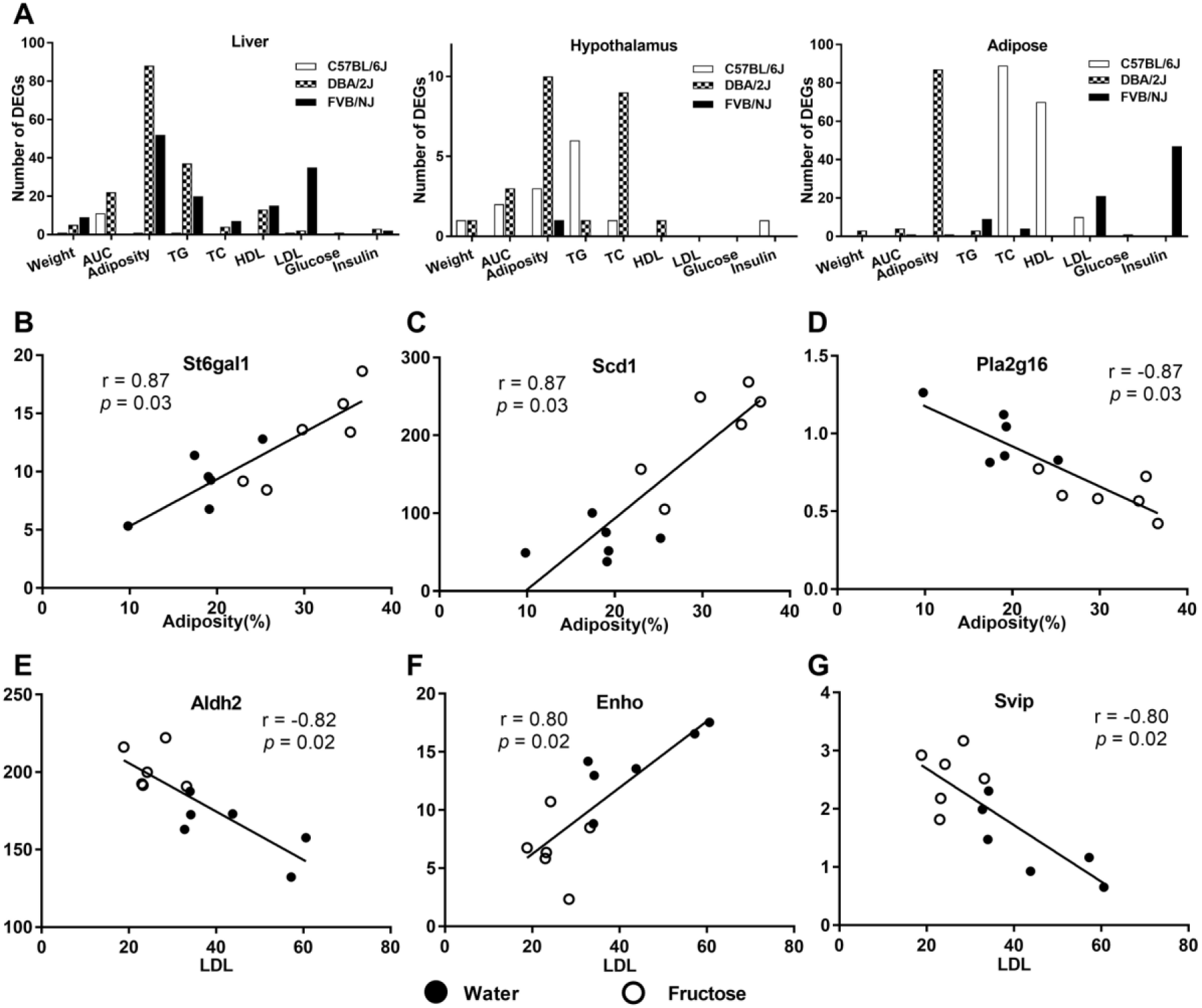
Relationship between DEGs and metabolic traits in mice. A: Numbers of strain-specific DEGs that are correlated with metabolic traits in our fructose study (*P* < 0.01 by Pearson correlation). B-D: Select examples of DEGs in liver correlating with adiposity in DBA mice. E-G: Select examples of DEGs in liver correlating with LDL in FVB mice.

To assess the association between the fructose DEGs with human cardiometabolic disorders (obesity, T2D, CAD, and lipids), we performed GWAS enrichment analysis using Mergeomics (42). As shown in Table 2, DBA liver and adipose DEGs showed strong enrichment for human GWAS signals for T2D and obesity traits; FVB DEGs from liver and adipose tissues are mostly related to lipid traits; B6 liver and adipose DEGs showed relevance to most cardiometabolic traits tested. These results suggest that the fructose-affected genes are relevant to human cardiometabolic diseases.

**Table 2.** Enrichment of human GWAS signals for metabolic syndrome related traits among strain- and tissue-specific DEGs based on Marker Set Enrichment Analysis (MSEA) in Mergeomics.

| Tissue of DEGs | Disease/trait | B6 | DBA | FVB |
| --- | --- | --- | --- | --- |
| Liver | CAD | **9.65E-05** | N.S | **2.85E-31** |
|  | T2D | **6.56E-08** | **4.02E-08** | N.S |
|  | HDL | **3.30E-05** | N.S | **8.90E-15** |
|  | LDL | 2.96E-03 | **3.77E-05** | N.S |
|  | TC | N.S | 2.57E-02 | 3.51E-03 |
|  | TG | **7.43E-12** | 1.93E-03 | 1.24E-03 |
|  | Fasting glucose | N.S | **4.60E-04** | 1.30E-03 |
|  | WCadjBMI | N.S | N.S | 1.54E-03 |
|  | HIPadjBMI | **2.37E-06** | **2.06E-09** | 2.77E-03 |
| Adipose | T2D | **1.72E-13** | N.S | N.S |
|  | HDL | **8.80E-41** | **1.25E-07** | 4.10E-02 |
|  | LDL | N.S | N.S | **8.09E-18** |
|  | TC | **9.41E-07** | N.S | N.S |
|  | TG | **1.87E-12** | 3.53E-03 | **8.12E-07** |
|  | Obesity | **4.89E-05** | **2.34E-10** | N.S |
|  | WCadjBMI | **3.58E-36** | N.S | **5.17E-07** |
|  | HIPadjBMI | **8.81E-23** | N.S | **1.17E-07** |
| Hypothalamus | HDL | 1.57E-02 | N.S | N.S |
|  | LDL | 1.57E-02 | N.S | N.S |
|  | TC | 9.64E-03 | N.S | N.S |
|  | TG | 2.36E-02 | N.S | N.S |
|  | Obesity | 2.41E-02 | N.S | N.S |

Note: Only traits and statistics with uncorrected MSEA enrichment $P < 0.05$ from at least one DEG list were shown. P values in bold passed Bonferroni-corrected $P < 0.05$ . CAD: coronary artery disease; T2D: type 2 diabetes; HDL: high density lipoprotein cholesterol; LDL: low density lipoprotein cholesterol; TC: total cholesterol; TG: triglycerides; WCadjBMI: BMI-adjusted waist circumference; HIPadjBMI: BMI-adjusted hip circumference. N.S.: not significant; $P > 0.05$ in MSEA.

## Discussion

Our multi-strain multi-tissue systems analysis shows that fructose consumption elicits differential phenotypic and transcriptomic profiles among mice of different genetic background. At the phenotypic level, DBA mice exhibited stronger vulnerability to alterations in body composition and glucose intolerance while B6 and FVB mice had stronger cholesterol alterations. We also report strain-specific regulatory genes and pathways in key metabolic tissues that may underlie the inter-individual phenotypic responses to fructose.

Our findings are in line with prior studies that demonstrated strain-specific responses to diets (6; 7) and offer a comprehensive account of the fructose-induced metabolic and molecular differences between strains. Interestingly, the patterns of phenotypic differences are not only strain-specific but also diet-specific. For example, among the three strains (B6, DBA, and FVB), B6 mice gained the most weight when fed a high fat high sucrose diet (7), whereas DBA mice gained the most weight when fed fructose diet. This diet-specific variability across strains indicates different adaptive abilities of distinct genetic background towards different diets. Our transcriptomic studies provide clues to the potential molecular underpinnings of the differential metabolic responses to fructose between mouse strains. In particular, we found very limited overlap between strains in the DEGs altered by fructose in individual tissues, and unique pathways represented by the strain-specific DEGs are biologically relevant to the distinct phenotypes in the corresponding mouse strains, as detailed below.

Liver-specific DEGs in the DBA mice were uniquely enriched for the PPAR signaling pathway, which agrees with results that DBA was the only strain to gain weight and fat mass and to show glucose intolerance in response to fructose. Indeed, PPAR gamma agonists are commonly used to improve insulin sensitivity and glucose intolerance and have influence on body weight (43). Network modeling revealed *Fgf21* to be a DBA-specific key regulator in the liver in response to fructose. FGF21 is a hormone primarily produced in the liver and holds promise as a potential therapy for obesity and glucose intolerance (44; 45). In our study *Fgf21* is significantly reduced by fructose in DBA mice, which agrees with the compromised glucose homeostasis and increased adiposity observed. Our *in vitro* fructose treatment experiments in primary hepatocytes confirmed that *Fgf21* is a direct and acute target of fructose in hepatocytes, particularly in DBA. There is also an interesting directional shift in *Fgf21* between the acute fructose response in primary hepatocytes where *Fgf21* immediately increases upon fructose treatment at early time points, whereas in our long-term (12-week) *in vivo* treatment study *Fgf21* is inhibited by fructose most significantly in DBA. Perhaps *Fgf21* is an early homeostatic regulator with a quick response to fructose upon initial exposure, but after long-term exposure, fructose or its metabolites or other downstream effectors inhibit *Fgf21*.

In contrast, B6 liver DEGs and top KDs are primarily cholesterol related genes, agreeing with the cholesterol-centric phenotypic responses of B6 to fructose. In FVB mice, fructose altered liver genes involved in vesicle-mediated transport pathways; the top KDs of FVB DEGs include *Itih3* and *Akr1d1;* DEGs *Aldh2, Enho*, and *Svip* are correlated with LDL. These genes are related to protein processing and metabolic pathways involving glutathione, cytochrome P450, and lipids, and may play roles in the unique decrease in plasma cholesterol and LDL levels in response to fructose in this strain.

In the hypothalamus, fructose induced the strongest transcriptomic changes in B6 mice compared to the other two strains, and our network analysis revealed extracellular matrix genes such as *Fmod* and *Bgn* as key drivers of the fructose-responsive DEGs. This is consistent with our previous study in which these genes were also identified as key regulators in the rat hypothalamus in response to fructose consumption (10). Both *Fmod* and *Bgn* knockout mice showed significantly altered cholesterol phenotypes (10), agreeing with the significant increases in these traits in B6 in response to fructose. The B6-specific increases in cholesterol species in response to fructose could also be related to dysregulation of liver key drivers related to cholesterol biosynthesis such as *Lss* as revealed in our network analysis. *Lss* encodes lanosterol synthase which converts (S)-2,3 oxidosqualene to lanosterol, a key intermediate in cholesterol biosynthesis, and there is growing interest in targeting this enzyme therapeutically to lower blood cholesterol (46).

In addition to the strain-specific DEGs and pathways, our transcriptomic analysis also revealed certain shared fructose DEGs across strains, which may imply robust targets of fructose regardless of genetic background. Some of these genes are involved in the response to xenobiotic stimulus (*Gstp1, Ephx1, Gstm1*) or organic cyclic compounds (*Abca1, Id3, Abat*), metabolic processes (*Abca1, Htatip2, Grhpr, Nudt7*), transcriptional regulation (*Ier5, Jun, Id3*), and immune modulation (*Cd200*). In particular, *Htatip2* modulates lipid metabolism through mediating the balance of lipid storage and oxidation (47). *Grhpr* encodes an enzyme involved in guiding the carbon flux to gluconeogenesis by converting hydroxypyruvate into D-glycerate (48). Many of the DEGs show correlations with metabolic traits in our mouse models and show significant over-representation of the candidate causal genes identified in GWAS of human cardiometabolic diseases.

To explore the gene regulatory mechanisms, we examined previously known transcription factors ChREBP and SREBP-1c (28–30). We found indirect evidence supporting their regulatory roles by observing changes in their target genes. Fructose metabolic enzymes such as *Khk* and *Aldob* also showed alterations but the direction of change and significance varied between strains in our 12-week fructose feeding study. More importantly, our data-driven network analysis revealed numerous additional strain-specific regulators of fructose activities in individual tissues, such as *Fgf21* (DBA), *Lss* (B6), and *Akr1d1* (FVB) in liver, BCAA genes (B6 and FVB) and *Dio2* (DBA) in adipose tissue, and extracellular matrix genes (B6) in hypothalamus. These context-specific regulatory genes identified here extend our knowledge about how fructose perturbs divergent pathways and triggers differential metabolic manifestations between individuals. Further investigations of these regulators, the genetic determinants of their context-dependent interactions with fructose or metabolites, and their downstream gene networks/pathways are warranted.

In conclusion, we found distinct metabolic phenotypes and molecular signatures in mice of different genetic background in response to fructose. These results provide important insight into how individuals differ in their response to fructose consumption by engaging specific gene regulatory mechanisms in both central and peripheral metabolic tissues. Future examination of additional mouse strains and tissues will help further dissect the inter-individual variability and tissue-specific genetic regulation of fructose response. Given the exponential rise of fructose consumption and concomitant increase in obesity and metabolic syndrome, our study provides key information for guiding personalized preventive and therapeutic strategies for diet-induced metabolic diseases.

## Author Contributions

G.Z., H.B., Z.Y., and K.C-K contributed to the experiments of the study. G.Z., J.H., X.Y. wrote the manuscript which was subsequently reviewed and revised by all the other authors. G.Z., M.B., Y.Z., L.S. performed the data analysis. X.Y., F. G-P. designed and supervised the study.

